# Wolbachia Infection Associated with Increased Recombination in Drosophila

**DOI:** 10.1101/238949

**Authors:** Nadia D. Singh

## Abstract

Wolbachia is maternally-transmitted endosymbiotic bacteria that infects a large diversity of arthropod and nematode hosts. Some strains of Wolbachia are parasitic, manipulating host reproduction to benefit themselves, while other strains of Wolbachia exhibit obligate or facultative mutualisms with their host. The effects of Wolbachia on its host are many, though primarily relate to host immune and reproductive function. Here we test the hypothesis that Wolbachia infection alters the frequency of homologous recombination during meiosis. We use *D. melanogaster* as a model system, and survey recombination in eight Wolbachia-infected and Wolbachia-uninfected strains, controlling for genotype. We measure recombination in two intervals of the genome. Our results indicate that Wolbachia infection is associated with increased recombination in one genomic interval and not the other. The effect of Wolbachia infection on recombination is thus heterogenous across the genome. Our data also indicate a reproductive benefit of Wolbachia infection; infected females show higher fecundity than their uninfected genotypic controls. Given the prevalence of Wolbachia infection in natural populations, our findings suggest that Wolbachia infection is likely to contribute to recombination rate and fecundity variation among individuals in nature.

## Introduction

Wolbachia is genus of Gram-negative bacteria that infects a wide variety of arthropod and nematode host species. As many as 40-66% of arthropods are infected with Wolbachia (Hilgenboecker et al. 2008; Zug and Hammerstein 2012) and in many cases, Wolbachia infection is observed at high frequencies in natural populations (e.g. Turelli and Hoffmann 1995). Wolbachia occupy the range of the parasite-mutualist spectrum in arthropods; there are myriad examples of Wolbachia behaving as a reproductive parasite as well as Wolbachia conferring fitness benefits to the host (for review see Zug and Hammerstein 2015). With respect to the former, Wolbachia have been shown to induce cytoplasmic incompatibility (e.g. Breeuwer and Werren 1990; Oneill and Karr 1990; Mercot et al. 1995; PerrotMinnot et al. 1996) and parthenogenesis (for review see Huigens and Stouthamer 2003; Engelstadter and Hurst 2009). Wolbachia infection also can manipulate its host to produce female-biased sex ratios in other ways including feminization and male-killing (for review see Kageyama et al. 2012).

In Drosophila, strains of Wolbachia span the parasite-mutualist spectrum as well. Male-killing strains of Wolbachia are present in a number of Drosophila species (Jaenike et al. 2003) (Hurst et al. 2000; Sheeley and McAllister 2009; Richardson et al. 2016), as are strains that induce cytoplasmic incompatibility (Hoffmann et al. 1986; Giordano et al. 1995; Bourtzis et al. 1996; Charlat et al. 2002; Richardson et al. 2016). However, in many cases Wolbachia does not appear to behave as a reproductive parasite in Drosophila. Strains that induce neither cytoplasmic incompatibility nor male-killing have been identified in a number of species (Hoffmann et al. 1996; Zabalou et al. 2004; Hamm et al. 2014).

In some cases Wolbachia infection appears to have a fitness benefit to the host. These fitness benefits largely fall into two categories: survival/reproduction and immune-defense. With respect to the former, several studies in *D. melanogaster* have revealed that Wolbachia-infected flies can enjoy enhanced survival and fecundity (Fry and Rand 2002; Fry et al. 2004; Serga et al. 2014). These fitness benefits could contribute to the high Wolbachia infection rates seen in natural populations of *D. melanogaster* (Serga et al. 2014). With regard to immune-related benefits, Wolbachia infection confers protection against viral infection in Drosophila, leading to reduced viral load and/or decreased mortality associated with viral infection (Hedges et al. 2008; Teixeira et al. 2008; Osborne et al. 2012; Chrostek et al. 2013; Stevanovic et al. 2015). Data on whether Wolbachia infection confers resistance or tolerance to bacterial infection are less clear. Although some studies have shown no Wolbachia-mediated antibacterial protection in *D. melanogaster* (Wong et al. 2011; Rottschaefer and Lazzaro 2012; Shokal et al. 2016), other data are suggestive a protective effect of Wolbachia infection against secondary infection with pathogenic bacteria (Unckless et al. 2015 though we note that the effects of host genotype cannot be ruled out). It has recently been suggested that the antibacterial protective effects of Wolbachia infection may be tied to whether the secondary infection is enteric or systemic, with protective properties evident in the case of enteric but not systemic infections (Gupta et al. 2017).

The demonstrated effects of Wolbachia on both reproductive and immune phenotypes in Drosophila led us to consider the potential effects of Wolbachia on meiotic recombination rate. We have recently shown that *D. melanogaster* females plastically increase their recombination fraction in response to pathogen attack, and that an active immune response appears key for recombinational response (Singh et al. 2015). Meiotic recombination occurs during oogenesis and Wolbachia colonize the female germline, so a potential link between Wolbachia infection and recombination seems intuitive. Moreover, recent work has shown that Wolbachia infection status is a significant predictor of recombination rate in *D. melanogaster* (Hunter et al. 2016a) though we hasten to note that this study, while suggestive of an effect, cannot rule out the confounding effects of genotype on recombination rate.

Here we test the extent to which Wolbachia infection alters the recombination fraction in *D. melanogaster* females. We survey recombination using a classical genetic approach in Wolbachia-infected and uninfected strains, controlling for genotype. We measured recombination in two genomic intervals—one autosomal and one X-linked. Our data indicate that recombination rates are significantly higher in Wolbachia-infected flies as compared to their genetically matched uninfected controls, but only for one of the two intervals surveyed. These data strengthen the link between infection and increased recombination, though in this case the effect is particularly interesting because the strain of Wolbachia in the current experiment is not considered to be pathogenic. We also find that although Wolbachia infection does not significantly affect the sex ratio of progeny produced, it does yield increased reproductive output for Wolbachia-infected females relative to uninfected females. Given the high frequencies of infection by Wolbachia in natural populations of *D. melanogaster,* our results suggest that variation in recombination rate and fecundity of *D. melanogaster* females in nature may be driven in part by Wolbachia.

## Materials and Methods

### Stocks and fly rearing

The eight wild-type lines used for this experiment were selected from the Drosophila Genetic Reference Panel (Mackay et al. 2012). These lines are: RAL149, RAL306, RAL321, RAL365, RAL790, RAL853, RAL879, and RAL897. All of these strains have standard chromosome arrangements. These line are naturally infected with *Wolbachia pipientis* (Huang et al. 2014). Genomic analysis indicates that the colonizing Wolbachia strain is a *wMel*-like strain (Richardson et al. 2012). *wMel* has been shown to induce cytoplasmic incompatibility, though the magnitude of the effect varies among studies (Bourtzis et al. 1996; Poinsot et al. 1998; McGraw et al. 2002; Reynolds and Hoffmann 2002; Yamada et al. 2007). We created Wolbachia-free versions of these eight strains. To cure the strains of Wolbachia infection, we raised flies on tetracycline-containing media for two consecutive generations. These flies were raised in 8 ounce (oz) bottles with a standard cornmeal/molasses media containing tetracycline (dissolved in ethanol) to a final concentration of 0.25 mg/ml media. After the second generation of tetracycline treatment, these strains (denoted, for example, RAL149^*w*-^) were raised on standard media for more than five generations before the experiment described below began.

We used doubly-marked strains for our estimation of recombination rate. The markers used to measure recombination on the X chromosome were *yellow* (*y*^1^) and *vermilion* (*v*^1^) (Bloomington Drosophila Stock Center #1509), which are 33 cM apart (Morgan and Bridges 1916). We integrated this doubly-marked X chromosome into the wild-type isogenic Samarkand genetic background (Lyman *et al.* 1996); this line abbreviated hereafter as ‘*y v*’. The markers on the chromosome 3R were *ebony* (*e*^4^) and *rough* (*ro*^1^) (Bloomington Drosophila Stock Center #496), which are 20.4 cM apart (Bridges and Morgan 1923); this line is abbreviated hereafter as ‘e *ro*.’ These markers and strains have been used extensively in our lab to estimate recombination frequency (Jackson et al. 2015; Singh et al. 2015; Hunter et al. 2016a; Hunter et al. 2016b).

### Wolbachia screen

Immediately prior to conducting these experiments, we confirmed the presence of Wolbachia infection in the standard RAL lines (denoted, for example, RAL149^*w*+^) and the absence of Wolbachia in the tetracycline-treated lines using a PCR-based assay with Wolbachia-specific primers. Four adult females were used per line to test for Wolbachia infection. Briefly, DNA was extracted from each female using a standard squish prep described by Greg Gloor and William Engels (personal communication). Each fly was crushed with a motorized pestle and subsequently immersed in a buffered solution (10 mM Tris-Cl pH 8.2, 1 mM EDTA, 25 mM NaCl, 200 μg/ml proteinase K). This was incubated at 37°C for 30 min and then at 95°C for 2 min to inactivate the proteinase K. We used Wolbachia-specific primers wspF and wspR (Jeyaprakash and Hoy 2000) to test for presence/absence of Wolbachia infection.

Amplifying conditions were as follows: 94°/3 min, 12 cycles of 94°/30 sec, 65°/30 sec, 72°/60 sec with the annealing temperature reduced by 1.0 degrees per cycle, followed by 25 cycles of 94°/30 sec, 55°/30 sec, 72°/60 sec. We included a final extension of 72°/7 min. All PCR reactions were 10 μl, and each contained 5 μl Qiagen 2X PCR Master-Mix, 0.25 μl of each 20 mM primer, 3.5 μl H_2_O, and 1 μl genomic DNA. All four tested females from each of the eight Wolbachia-infected lines showed evidence of Wolbachia infection using this assay, while none of the females from the tetracycline-treated lines showed any evidence for Wolbachia infection.

### Experimental crosses

To assay recombination rate variation in the experimental lines, we used a classic two-step backcrossing scheme. All crosses were executed at 25° C with a 12:12 hour light:dark cycle on standard media using virgin females aged roughly 24 hours. For the first cross, ten virgin females from each experimental evolution line were crossed to ten *e ro* (or *y v)* males in eight oz bottles. Males and females were allowed to mate for five days, after which all adults were cleared from the vials. F_1_ females resulting from this cross are doubly heterozygous; these females are the individuals in which recombination is occurring. To uncover these recombination events we backcrossed F_1_ females to doubly-marked males. For this second cross, twenty heterozygous virgin females were collected and backcrossed to twenty doubly-marked males in 8 oz bottles. Males and females were allowed to mate for five days, after which all adults were cleared from the bottles. After eighteen days, BC_1_ progeny were collected and scored for sex and for visible phenotypes. Recombinant progeny were then identified as having only one visible marker (*e*+ or +*ro* in the case of crosses involving the *e ro* double mutant, or *y*+ or +*v* in the crosses involving the *y v* double mutant). Ten to fifteen replicate (second) crosses were set up for each strain. For each replicate, recombination rate was estimated by taking the ratio of recombinant progeny to the total number of progeny. Double crossovers cannot be recovered with this assay, so our estimates of recombination frequency are likely to be biased downwards slightly.

### Statistical analyses

All statistics were conducted using JMPPro v13.0. To test for factors associated with variation in recombination fraction, we used a generalized linear model with a binomial distribution and logit link function on the proportion of progeny that is recombinant. We treated each offspring as a realization of a binomial process (either recombinant or nonrecombinant), summarized the data for a given vial by the number of recombinants and the number of trials (total number of progeny per vial), and tested for an effect of line and Wolbachia status plus the interaction of line and Wolbachia status. The full model is as follows:

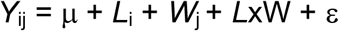

for: *i* = 1…8, *j* = 1…2

*Y* represents the proportion of progeny that is recombinant, μ represents the mean of regression, and ε represents the error. *L* denotes strain, *W* denotes Wolbachia infection status, and *L*x*W* denotes the interaction of line and Wolbachia infection status. All of these are modeled as fixed effects.

To test for factors associated with variation in the sex ratio, we used the same generalized linear model framework with a binomial distribution and logit link function on the proportion of total number of progeny that is male. For each interval, the model is the same as is described above, except that *Y* represents the proportion of progeny that is male. We note that this analysis is independent of recombination frequency.

To test for factors associated with variation in reproductive output, we used an ANOVA framework. The ANOVA followed the form of *Y* = *μ* + *L* + W + *ϵ*, for each interval assayed where *Y* is reproductive output, *μ* is the overall mean, *L* is the fixed effect of line, *W* is the fixed effect of Wolbachia status, and *ϵ* is the residual.

## Results

### Viability effects of markers on recombination rate estimation

In total, 76,211 BC1 progeny were scored for recombination for the *e ro* interval, and 79,447 flies were scored for recombination in the *y v* interval. For the *e ro* interval, the number of progeny per bottle ranged from 123-732, with an average of 476 BC1 progeny per vial. For the *y v* interval, the number of BC1 progeny ranged from 230-630, with an average of 462 progeny per bottle.

To test for deviations from expected ratios of phenotype classes, we performed *G*-tests for goodness of fit for all crosses for the following ratios: males versus females, wild-type flies versus *e ro* (or *y v*) flies and finally, *e* + flies versus + *ro* flies (or *y* + versus + *v* flies). The null hypothesis for each comparison is a 1:1 ratio of phenotype classes. We tested for such deviations within each bottle.

Examining the ratio of males to females for BC1 progeny associated with the *e ro* interval, 9 of 160 crosses (6%) show a significant deviation from the expected 1:1 ratio (*P* < 0.05, *G*-test). Of these 9 crosses, six show a female-bias in the progeny count. None of these deviations remain significant after using a Bonferroni-correction for multiple tests.

Comparing the number of wild-type to *e ro* flies, 30 of 160 crosses (19%) deviate significantly from expectation. In 28 of those crosses, there is an excess of wild-type flies relative to the double mutant. The ratio of wild-type to double mutant ranges from 1.2 to 1.4, with an average of 1.3. One of these crosses shows a significant deviation from expectation after correcting for multiple testing (Bonferroni-corrected *P* = 0.03, G-test).

With respect to the ratio of *e*+ / +*ro* flies, seven crosses (4%) show a significant departure from the expected 1:1 ratio. Five of these show an excess of +*ro* flies, while two show an excess of *e*+ flies. None of these departures remain significant after using a Bonferroni-correction for multiple tests.

Comparing total females to total males for the progeny associated with the *y v* interval, 20 of 172 (22%) crosses show significant deviations from the expected 1:1 ratio (*P* < 0.05, *G*-test). A relative excess of females is observed in 15 of those crosses. The female biased crosses show M/F ratios ranging from 0.70-0.83, with an average of 0.81. Only one of these deviations remains significant after using a Bonferroni-correction for multiple tests (Bonferroni-corrected *P* = 0.02, G-test).

With respect to wild-type versus *y v* flies, 21 of 172 (12%) crosses show a significant deviation from the expected 1:1 ratio (*P* < 0.05, G-test), and in all but six of these cases these crosses yield a relative excess of wild-type flies. The ratio of wild-type to double mutant in those wild-type biased crosses ranges from 1.2-1.8, with an average of 1.3. This is consistent with a mild viability defect associated with the visible markers. However, only one of these deviations remains significant after using a Bonferroni-correction for multiple tests (Bonferroni-corrected *P* = 0.0008, G-test).

Finally, 12 of 173 (7%) lines show significantly different numbers of *y*+ flies versus +*v* flies (*P* < 0.05, G-test), with 10 of those 12 crosses showing an excess of +v flies. The *y*+/+*v* ratio for these 10 lines ranges from 0.58-0.71, with an average of 0.65 across the 10 crosses. None of these deviations remain significant after using a Bonferroni-correction for multiple tests.

Although the number of crosses with significant deviations from null expectation is small relative to the total number of crosses, our data are nonetheless indicative of a mild viability defect associated with our marked chromosome. However, these skewed ratios do not appear depend on Wolbachia infection status. Fitting a generalized linear model with a binomial distribution and logit link function on the proportion of non-recombinant progeny that is wild-type shows no significant effect of Wolbachia status (*P* = 0.07, χ^2^ test (N = 172, df = 7)) for the *y v* crosses. The same result is found with the *e ro* crosses (*P* = 0.42, χ^2^ test (N = 160, df = 7)). We thus believe that the small viability defects associated with the doubly marked chromosome are not systematically biasing the estimates of recombination with respect to Wolbachia status in this experiment.

### Effects of Wolbachia infection status on recombination

We used a logistic regression model to identify factors significantly contributing to variation in recombination fraction observed in the current experiment. Note that because we are assaying recombination in heterozygous females (see Materials and Methods), we can only detect dominant genetic effects. The logistic regression model indicates that both genotype and Wolbachia infection status significantly contribute to variation in recombination rate in the *y v* interval (*P* < 0.0001, both factors, χ^2^ test (N = 172, df = 7 for genotype, df = 1 for Wolbachia status)). Average recombination values for each strain are presented in Figure 1, and these data illustrate that recombination fraction increases with Wolbachia infection. There is no significant interaction effect between Wolbachia status and genotype (*P* = 0.54, χ^2^ test (N = 172, df = 7)).

Different results are found for the *e ro* interval. Although genotype significantly contributes to variation in recombination fraction (*P* < 0.0001, χ^2^ test (N = 160, df = 7)), Wolbachia status does not (*P* = 0.31, χ^2^ test (N = 160, df = 1)). The lack of consistent effect of Wolbachia status on recombination fraction is echoed in Figure 1. There is a marginally significant interaction effect between Wolbachia status and genotype on recombination fraction (*P* =0.04, χ^2^ test (N = 160, df = 7)). However, the distributions of recombination fractions for a given genotype with Wolbachia and without Wolbachia do not differ significantly in any of the eight comparisons (*P* > 0.08, all tests, Wilcoxon rank test).

**Figure 1:**
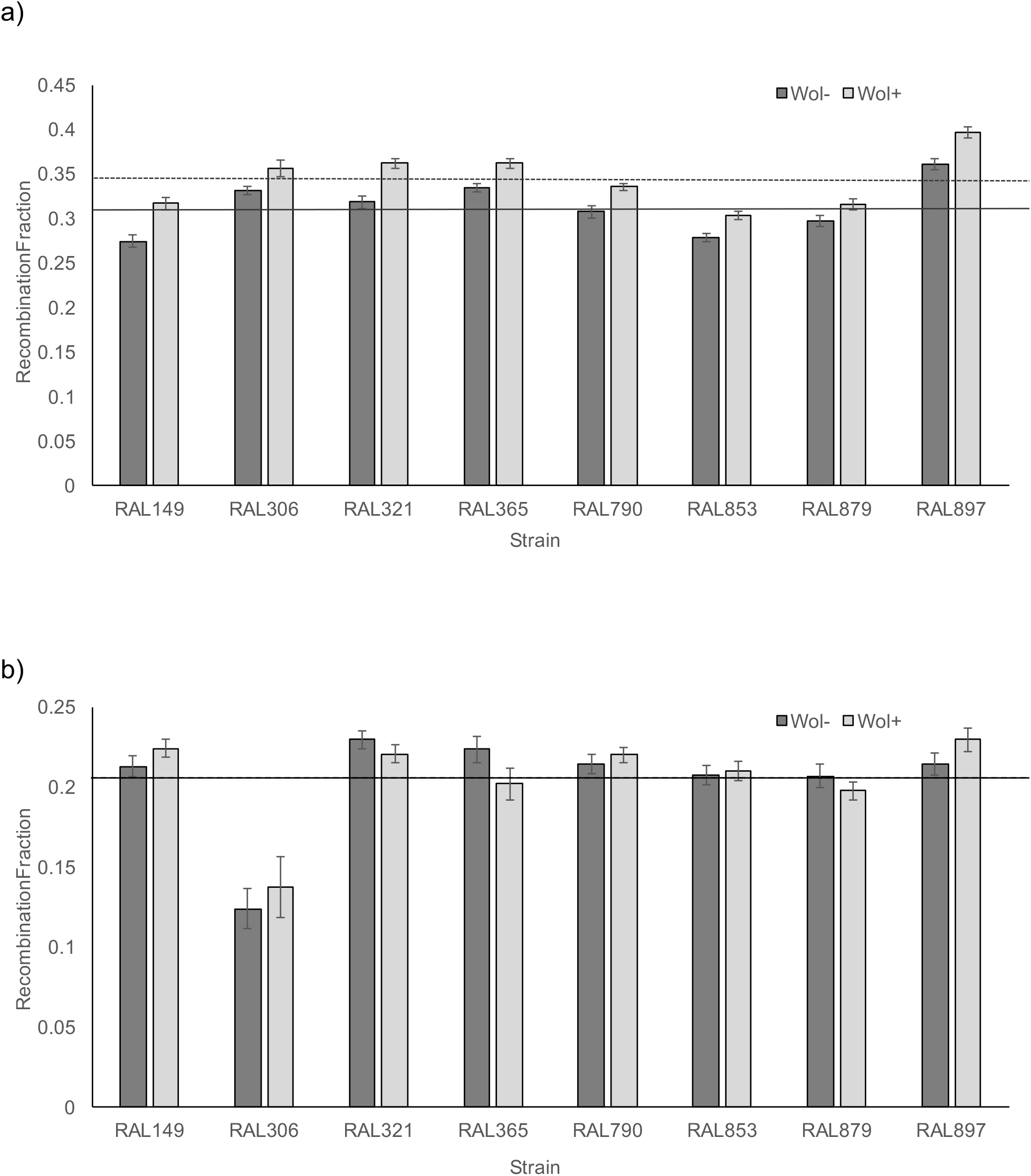
Mean recombination fraction between a) *yellow* and *vermillion* and b) *ebony* and *rough* interval as a function of genetic background and Wolbachia infection status. Dark grey bars are Wolbachia-free lines and the light grey bars are the Wolbachia-infected counterparts. Error bars denote standard error. Average recombination rate (across lines) is depicted for Wolbachia-infected (dashed line) and uninfected lines (solid line).

### Effects of Wolbachia infection status on other phenotypes

Given the known effects of Wolbachia infection on host biology, we tested for effects of Wolbachia infection on two additional phenotypes: reproductive output and sex ratio. To examine factors associated with variation in reproductive output, we used an ANOVA framework. For the crosses involving *y v* flies, both genotype and Wolbachia status significantly affect reproductive output (as measured by the number of progeny in a replicate) (*P*_Genotype_ = 0.03, *P*_Wolbachia_ = 0.01, respectively). Figure 2 illustrates that reproductive output generally increases with Wolbachia infection. There is no significant interaction between genotype and Wolbachia status (*P* = 0.63). For the crosses involving *e ro* flies, genotype and Wolbachia status again contribute to reproductive output (*P*_Genotype_ < 0.0001, *P*_Wolbachia_ = 0.003, respectively); Wolbachia infection results in an increase in reproductive output (Figure 2b). For these crosses, there is also a significant interaction effect between Wolbachia status and genotype (*P* = 0.03), with Wolbachia status impacting reproductive output in some strains more strongly than others (Figure 2b).

**Figure 2:**
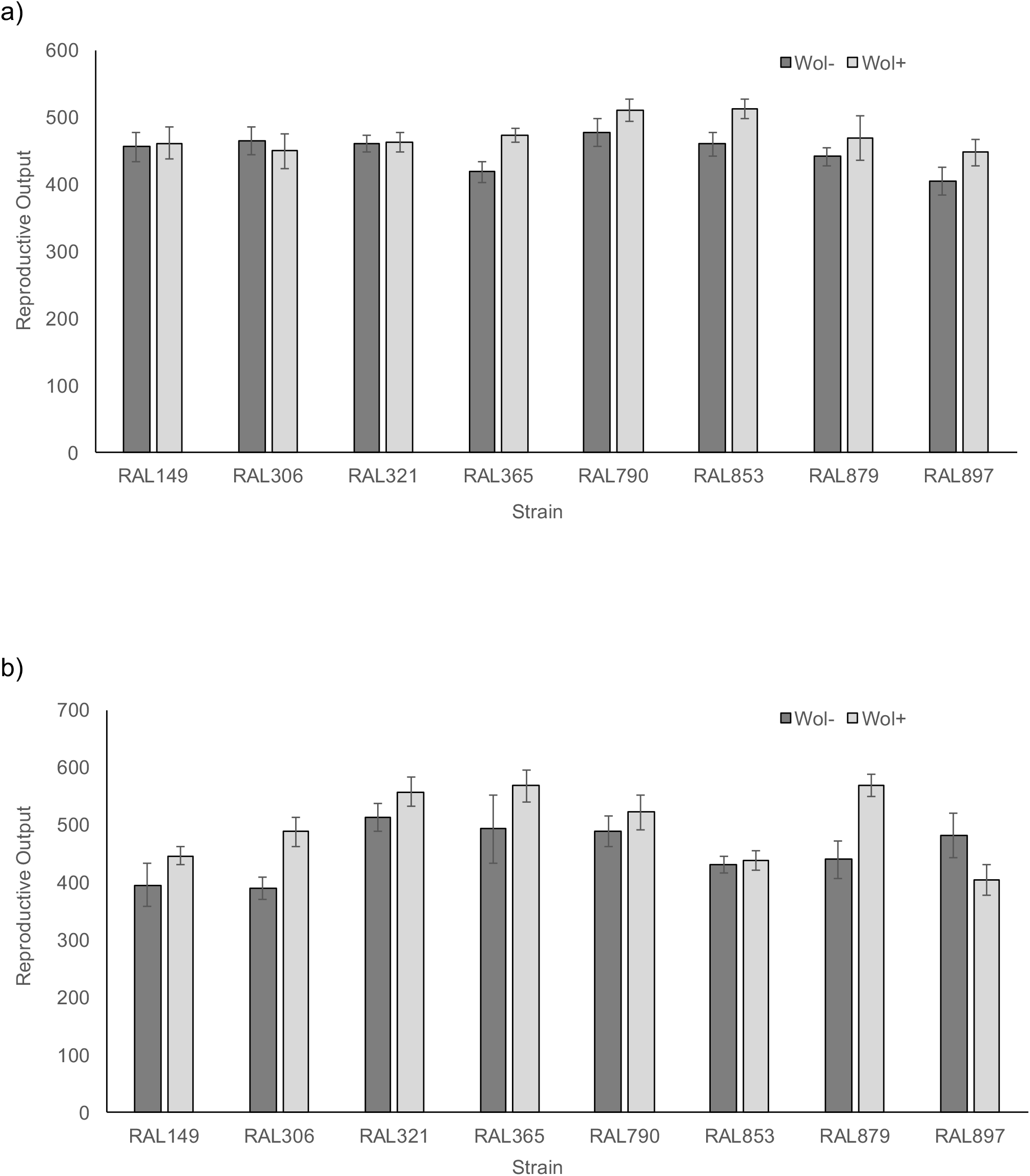
Mean reproductive output for crosses measuring recombination in the a) *yellow* and *vermillion* and b) *ebony* and *rough* interval as a function of genetic background and Wolbachia infection status. Dark grey bars are Wolbachia-free lines and the light grey bars are the Wolbachia-infected counterparts. Error bars denote standard error. Average recombination rate (across lines) is depicted for Wolbachia-infected (dashed line) and uninfected lines (solid line), though they are not both visible because they are so close together (0.204(uninfected) versus 0.205(infected)).

To test for factors affecting the sex ratio, we used a logistic regression (see Materials and Methods). For the crosses involving *y v* flies, there are no significant effects of genotype, Wolbachia status, or the interaction between genotype and Wolbachia status on the ratio of males to females (*P* = 0.08, 0.55, and 0.48, respectively). Progeny resulting from backcrosses with *e ro* flies show the same result, with no significant contribution of genotype, Wolbachia status, or the interaction between genotype and Wolbachia status to the sex ratio (*P* = 0.10, 0.19, and 0.16, respectively)

## Discussion

Here we show that infection with Wolbachia significantly increases the recombination fraction for an X-linked genomic interval but not an autosomal region (Figure 1). This recapitulates one of the motivating findings that Wolbachia infection was a significant predictor of recombination level for the X-linked *y v region* but not the autosomal *e ro* region (Hunter et al. 2016a). However, the effects of Wolbachia infection in that study were conflated with genotype, so we could not exclude genotype as driving the observed variation in recombination between Wolbachia-infected and Wolbachia-free strains. A more direct test of the effect of Wolbachia status on recombination rates would control for genotype; previous work with this experimental framework revealed that Wolbachia infection had no detectable effect of levels of crossing over in *D. melanogaster* (Serga et al. 2010). The discrepancy between that finding and the current study may be due to the fact that different genotypes were used in the two studies. In addition, different genomic intervals were surveyed in the two studies (though we note that our X-linked locus encompasses that utilized in the initial study). Indeed, the degree to which recombination is plastic varies among genotypes and among loci in *D. melanogaster* (Hunter et al. 2016b).

That one genomic interval shows a plastic response to Wolbachia infection while another does not is also consistent with previous reports indicating variation in recombination plasticity across intervals in *D. melanogaster* (Hunter et al. 2016b) and *D. pseudoobscura* (Stevison et al. 2017). Interestingly, the *e ro* interval surveyed here has previously been shown to exhibit a plastic increase in recombination following infection with pathogenic bacteria (Singh et al. 2015) and heat shock (Jackson et al. 2015), but no plastic response to maternal age (Hunter et al. 2016b). This suggests that genomic intervals may exhibit phenotypic plasticity in recombination in response to some environmental cues but not others. The *y v* interval surveyed in the current study, showing an increase in recombination associated with Wolbachia infection also shows an increase in recombination with maternal age (Hunter et al. 2016b). It remains to be determined what causes some loci to exhibit plastic recombination under certain conditions.

That Wolbachia affects recombination in an X-linked interval but not an autosomal one is potentially of interest. Because Wolbachia has been shown to affect the sex ratio in many systems, there may be some interaction between Wolbachia and the X chromosome. Surveying recombination in additional sex-linked and autosomal regions is required to determine whether the observations in the current study are reflective of broader patterns. Future work could also include cytological assays to determine whether the X undergoes more meiotic double-stranded breaks in Wolbachia-infected versus uninfected flies, and if autosomal breaks change in frequency in a Wolbachia-dependent way.

Recombination plasticity associated with bacterial infection has been previously shown in *D. melanogaster* as well (Singh et al. 2015). In contrast to the pathogenic bacteria used in that experiment (*Serratia marcescens* and *Providencia rettgeri),* the *wMel* strain of *Wolbachia pipentis* used in the current experiment is viewed more as a mutualistic endosymbiont rather than a pathogenic bacterium. However, the plastic increase in recombination associated with bacterial infection with pathogenic bacteria appeared to be driven in part by an active immune response (Singh et al. 2015). Several studies have indicated that the *D. melanogaster* transcriptional profile significantly differs between Wolbachia-infected and Wolbachia-free flies. In vitro examination of S2 cells challenged with Wolbachia showed upregulation of several immune genes including *Toll, Imd,* and five antimicrobial peptides (Xi et al. 2008). Similarly, larval testes show significant upregulation of nine immune-related genes in Wolbachia-infected males compared with uninfected controls (Zheng et al. 2011). However, other results show no effect of Wolbachia alone on regulation of immune genes (Bourtzis et al. 2000; Wong et al. 2011; Shokal et al. 2016). A key difference among the studies is the type of Wolbachia strains used for infection, which could underlie the differences in immune response observed among studies. It appears as though those studies showing a significant effect of Wolbachia on immune function were limited to those studying the effects of Wolbachia on hosts that were naturally infected with a Wolbachia strain that was different from the strain used for experimentation (for review see Zug and Hammerstein 2015). Here we leveraged naturally-infected hosts, which could result a lack of immune-system activation in the current experiment, though testing this explicitly with comparative transcriptomics of naturally Wolbachia-infected and uninfected hosts is an avenue of future exploration.

In addition to potentially affecting immune signaling, Wolbachia infection has been associated with key changes in germline function. This is of importance to our study given its focus on meiotic recombination, which occurs in the female germline. *Drosophila mauritiana* females show both an increase in egg production as well as an increase in mitotic activity of germline stem cells associated with Wolbachia infection (Fast et al. 2011). In *D. melanogaster,* Wolbachia infection reduces maternal transmission of the endogenous retrovirus *gypsy* (Touret et al. 2014). The presence of Wolbachia also significantly affects protein abundance in *D. melanogaster* ovaries (Christensen et al. 2016). Wolbachia infection also increases the frequency of apoptosis in oogenesis in both mosquitoes (Almeida and Suesdek 2017) and *D. melanogaster* (Zhukova and Kiseleva 2012). This latter finding is particularly intriguing; though we note that a virulent strain of Wolbachia was used (wMelPop), the stage of oogenesis that was examined and showed higher levels of apoptosis associated with Wolbachia infection was region 2a/2b of the germarium. This is precisely where double strand breaks are initiated and resolved (Jang et al. 2003; Staeva-Vieira et al. 2003; Gorski et al. 2004). This suggests that Wolbachia infection has the capacity to affect host reproductive phenotypes in exactly the region of the germarium in which crossovers are formed. Another mechanistic link between meiotic recombination and Wolbachia infection is found in the gene *lola. lola* is one of five high confidence candidate genes at which allelic variation is significantly associated with population-level variation in recombination rate (Hunter et al. 2016a), and *lola* also shows significant downregulation in larval testes in the presence of Wolbachia. Disruptions in *lola* yield increases in recombination rate (Hunter et al. 2016a) which is consistent with the predicted direction of the effect given the effect of Wolbachia on *lola* expression (Zheng et al. 2011) and its effect on recombination observed in the current study.

Consistent with its effect of on the female germline, our data further indicate that Wolbachia infection significantly increases reproductive output (Figure 2). This echoes what has been shown in *D. simulans* (Weeks et al. 2007), *D. mauritiana* (Fast et al. 2011) and *D. suzukii* (Mazzetto et al. 2015 though see also Hamm et al. 2014). Consistent with this, in *D. melanogaster,* females infected with mutualistic strain *wMel* (denoted wDm in some early literature) generally show higher reproductive output than Wolbachia-uninfected flies (Fry et al. 2004; Serga et al. 2014). Moreover, fecundity of *D. melanogaster* females infected with the more rare strain wMelCS show reduced egg production relative to those flies infected with *wMel* (Serga et al. 2014). This gives some credence to the idea that infections with *wMel* benefit the host, and that these benefits may have evolved from the stability of the host-endosymbiont relationship given vertical transmission.

It should be noted that recent data indicate that the Wolbachia-associated fitness benefits realized through female fecundity can be context-dependent. Diet, for instance, plays a major role in mediating the effect of Wolbachia on female reproductive output. Wolbachia-infected flies reared on protein-rich diets or high-or low-iron diets show increased fecundity relative to their Wolbachia-free counterparts (Brownlie et al. 2009; Ponton et al. 2015). This indicates that nutrient availability can mediate the effects of Wolbachia infection on its host, and highlights the complex nature of the interactions among host, endosymbiont, and the environment.

Unlike reproductive output, the sex ratio of progeny produced did not differ systematically between Wolbachia-infected and uninfected flies in the current study. This mirrors previous results in *D. melanogaster* (with or without wMel), which also showed no significant difference in sex ratio associated with Wolbachia status (O’Shea and Singh 2015; Ponton et al. 2015). Although Wolbachia strains that manipulate the sex ratio clearly exist in other Drosophila species (Hurst et al. 2000; Dyer et al. 2005; Sheeley and McAllister 2009; Richardson et al. 2016), wMel does not appear to have a feminizing effect in *D. melanogaster.* However, because as noted above the environment can shape the effects of Wolbachia on its host, it remains possible that wMel could shift the sex ratio in *D. melanogaster* under different environmental conditions.

## Conclusions

Recombination frequency is phenotypically plastic in a number of organisms. This is perhaps best studied in *Drosophila melanogaster,* where the first reports of plastic recombination were published nearly a century ago. Here we test the hypothesis that Wolbachia infection increases recombination in *D. melanogaster.* This hypothesis was motivated by two previous studies, one showing that infection with pathogenic bacteria yields a plastic increase in recombination, and one showing that Wolbachia status was significantly associated with population level-variation in recombination. This latter study did not inform the extent to which Wolbachia infection could yield increased recombination because the effects of Wolbachia could not be separated from effects of genotype. By surveying recombination at two loci in eight wild-type strains of *D. melanogaster*, we show that infection with Wolbachia is associated with increased recombination in one genomic interval but not another interval. It is not known whether this effect is modulated by the immune system--this is a topic of future work. The factors underlying the genomic heterogeneity in the plastic recombinational response are unknown as well. We further find that reproductive output is increased with Wolbachia infection, indicating a benefit of infection to the host. Given that Wolbachia are materally-transmitted, increased reproductive output of the fly host boosts not only host reproductive fitness, but the fitness of Wolbachia as well.

## Acknowledgments

The author gratefully acknowledges Rob Unckless for feedback on the manuscript. This work was supported by National Science Foundation grant MCB-1412813 to N. D. S.

